# Graph analysis of structural brain networks in Alzheimer’s disease

**DOI:** 10.1101/050708

**Authors:** Majnu John, Toshikazu Ikuta, Janina Ferbinteanu

## Abstract

**Background:** Changes in brain connectivity in patients with early Alzheimer’s disease (AD) have been investigated using graph analysis. However, these studies were based on small data sets, explored a limited range of network parameters, and did not focus on more restricted sub-networks, where neurodegenerative processes may introduce more prominent alterations.

**Methods:** In this study, we constructed structural brain networks out of 87 regions by using data from 135 healthy elders and 100 early AD patients selected from the Open Access Series of Imaging Studies (OASIS) database. We evaluated the graph properties of these networks by investigating metrics of network efficiency, small world properties, segregation, product measures of complexity, and entropy. Because degenerative processes take place at different rates in different brain areas, analysis restricted to sub-networks may reveal changes otherwise undetected. Therefore, we first analyzed the graph properties of a network encompassing all brain areas considered together, and then repeated the analysis after dividing the brain areas into two sub-networks constructed by applying a clustering algorithm.

**Results:** At the level of large scale network, the analysis did not reveal differences between AD patients and controls. In contrast, the same analysis performed on the two sub-networks revealed modifications accompanying AD. Changes in small world properties suggested that the ability to engage concomitantly in integration and segregation of information diminished with AD in the sub-network containing the areas of medial temporal lobe known to be heaviest and earliest affected. In contrast, we found that the second network showed an increase in small world propensity, a novel metric that unbiasedly quantifies small world structure. Complexity and entropy measures indicated that the intricacy of connection patterns and structural diversity decreased in both sub-networks.

**Conclusions:** These results show that neurodegenerative processes impact volumetric networks in a non-global fashion. Our findings provide new quantitative insights into topological principles of structural brain networks and their modifications during early stages of Alzheimer’s disease.

## INTRODUCTION

Alzheimer’s disease (AD) is a progressive neurodegenerative process that interferes with multiple brain functions including memory, language, attention, and perceptual skills. As the life span of the population is increasing, so are the prevalence of this illness and its costs to society. Early detection of AD is important because it can lead to increased efficacy of treatment. Currently, growing amounts of evidence from neuropathological, electrophysiological and neuroimaging studies support the hypothesis that AD includes a disconnection syndrome^1^ generated by a breakdown of the organized structure and function of multiple brain areas even in the early stages of the disease^2^. One useful source of information for early AD diagnosis is structural MRI, which can reveal abnormalities in a wide range of brain areas. Multiple studies have found structural MRI (sMRI) a useful diagnostic tools that can contribute to detecting AD-related modifications before the development of clinical symptoms^3-5^^6 7^. A different way in which sMRI can assist in early AD diagnosis is by providing data sets for a distinct group of recently developed analytical procedures involving advanced mathematical and statistical methods such as machine learning procedures^8-11^ and graph theory, the formal study of networks^12 13 14^.

Graph theoretical methods provide a powerful approach for quantifying the organization of network connectivity using brain anatomical features including gray matter volume, cortical thickness, surface area, and white matter pathways between gray matter regions^14 15 16 17 18 19 20^. When applying graph theory to sMRI data, the outcome is a network whose *nodes* or *vertices* are represented by brain regions or voxels defined by a predetermined parcellation scheme, while *edges* are represented by inter-individual data associations of these regions evaluated, for example, as the strength of correlation between regional measurements. (It should be mentioned that edges of structural networks are not always represented by correlations between regional measurements, but by the number or density of white matter tracts connecting regions; however, this was not the course our analysis took). Structural network analysis can be employed to characterize the properties of large scale brain circuits^21^, and changes in the structure of a network, such as anatomical alterations produced by the neurodegenerative processes in AD, modify the network’s properties, and likely its function. Accurate evaluation of these modifications through assessment of morphometric correlations between brain areas may provide novel insights into the modifications introduced by AD and suggest novel treatment avenues^22^.

Several previous studies of AD from the graph theory perspective reported aberrations in small world properties (ability to simultaneously integrate and segregate information processing) of whole brain structural networks generated by modifications in cortical thickness measurements^14 15 23^; decreased functional connectivity^12 13^; and high amyloid-β deposition in the locations of cortical *hubs* (highly connected nodes, which correspond to brain areas with high input/output in this type of analysis^24^. The latter finding is consistent with the possibility that hubs, while acting as critical way stations for information processing, may also augment the underlying pathological cascade in AD as neurodegeneration may progress preferentially along neural pathways (but see^25^). Although these studies represent significant progress in the study of AD, three out of the four structural studies cited above were limited by low sample sizes (less than 30 subjects/group). Thus, we aimed to base our investigation on a large group size and. To this purpose, we used 235 subjects selected from the Open Access Series of Imaging Studies. Second, the above studies focused mainly on only one class of network parameters, the small-world property, which quantifies the ability of a network to simultaneously perform functional integration and segregation. However, brain modifications in early AD may involve other changes in network parameters beside small world properties, which may provide additional insight into pathological processes. Therefore, in the current study we additionally used entropy, complexity, and efficiency to assess structural alterations of gray matter volume in AD patients. Third, in early AD neurodegeneration affects distinct brain areas to various degrees and consequently, modifications in brain networks are not uniform. To evaluate whether this heterogeneity can be captured by graph analysis, we investigated structural changes first in one large network, constituted from all the brain areas under consideration, and a second time after we clustered the brain areas using an algorithm that iteratively removes edges from the network to divide it into distinct groups^26^.

## METHODS

### Subjects

MRI, demographic and clinical data were obtained from the Open Access Series of Imaging Studies (OASIS) database (http://www.oasis-brains.org). The details of data acquisition were previously described^27^.The current analysis included two hundred and thirty-five subjects whose clinical data were available. In this sample, subjects with dementia are only those with AD-caused dementia^27^. Subjects who had a Clinical Dementia Rating (CDR) score^28 29^ equal to zero were classified as healthy elders (n = 135), and subjects who had CDR score 0.5 or above were treated as early stage AD patients (n= 100); 70 patients had a CDR score of 0.5, 28 a score of 1, and 2 a score of 2. The median age in the control group was 71 and age ranged from 33 to 94. The median age in the patients group was 77 with a range from 62 to 96. Mean ages for the controls and patients were 69.1 years and 76.8 years, respectively. 72% of the healthy controls and 59% of the early stage AD’s were females. Median Mini Mental State Exam (MMSE) score^30^, which measures the cognitive function, was 29 (range: 25 to 30) in controls and 26 (range: 14 to 30) in patients. Because age and gender influence correlation between regional volumes, we removed the effects of these two factors by regressing them on the structural volume of each region to obtain studentized residuals, which were then used for all further analyses.

### Volumetric MRI Data Processing

Structural T1 MRI data (256x256x160 voxels of 1mm^3^) were processed by FreeSurfer (version 5.3.0) in which volumetric parcellation was processed by recon-all version 1.379 with *–all* option. From cortical *aparc* parcellation and subcortical *aseg* segmentation outputs^31,32^, eighty seven cortical gray matter and subcortical regions including brain-stem and cerebellar cortex were incorporated in the analysis. All regions other than brain stem were separately and independently treated for left and right volumes. All brain regions are listed in Table 1.

**Table 1.** **Clustering of the brain regions.**

| **Cluster 1** | **Cluster 2** |
| --- | --- |
| R. supramarginal gyrus | L. caudate |
| L. and R. transverse temporal gyrus | R. thalamus |
| L. and R. paracentral gyrus | L. caudal middle frontal cortex |
| R. cuneus | L. cuneus |
| L. thalamus | L. and R. fusiform gyrus |
| brain stem | L. and R. middle temporal cortex |
| L. and R. pericalcarine cortex | R. medial orbitofrontal cortex |
| L. rostral anterior cingulate cortex | L. and R. rostral middle frontal cortex |
| L. and R. pars triangularis | L. and R. precuneus |
| L. and R. entorhinal cortex | L. precentral gyrus |
| L. and R. ventral diencephalon | L and R. superior temporal cortex |
| L. and R. pallidum | R. posterior cingulate cortex |
| L. and R. caudal anterior cingulate cortex | R. insula |
| L. and R. putamen | R. pars opercularis |
| L. and R. hippocampus | R. banks of superior temporal sulcus |
| L. and R. cerebellar cortex | L. and R. inferior temporal cortex |
| L. and R. amygdala | L. and R. lateral occipital cortex |
| L. and R. accumbens area | L and R. postcentral gyrus |
| L. posterior cingulate cortex | L. and R. superior parietal cortex |
| L. and R. lingual gyrus | R. rostral anterior cingulate cortex |
| L. insula | R. caudal middle frontal cortex |
| L. and R. pars orbitalis | L. and R. superior frontal cortex |
| L. and R. isthmus cingulate cortex | L. and R. lateral orbitofrontal |
| L. and R. parahippocampal cortex | L. and R. inferior parietal cortex |
| L. banks of superior temporal sulcus | L. supramarginal gyrus |
| L. pars opercularis |  |
| L. and R. frontal pole |  |
| L. and R. temporal pole |  |
| R. caudate |  |
| L. medial orbitofrontal cortex |  |
| R. precentral gyrus |  |

### Brain networks as graphs

Graphs are mathematical constructs of relationships among various objects. In graph models, the objects of interest are modeled as *nodes* (or vertices) and the relationships among the objects are modeled using *edges* (or lines) connecting the vertices. The brain can be conceived as a multitude of regions forming a complex network whose properties can be studied based on their graph theoretical properties. In our analysis, the nodes were gray matter volumes of 87 brain regions of interest. One way to model the relationships between the regions is by considering the correlations of the corresponding regional volumes. This could, in turn, lead to two types of graph models: a weighted graph model or a binary graph model. In either of these models, an edge between a pair of nodes represents the relationship that we are trying to capture between these nodes. In a binary graph model, for a pair of nodes, an edge simply exists or not, depending on whether the correlation between the volumes of gray matter in the corresponding brain areas passes a certain threshold. For example, to construct a binary graph when the relationship between a pair of nodes is based on correlation we may use a threshold value of 0.5, and declare that an edge exists between the pair of nodes if and only if the correlation is above 0.5. If we do this for all pairs of nodes, we will obtain the binary graph. In contrast, in a weighted graph, if an edge exists between a pair of nodes, it has also a numerical weight associated with it. In this case, if we proceed as above but also assign the actual correlation between the pair of nodes as the weight of the edge, then the result is a weighted graph. In mathematical notation, any binary graph with n nodes could be represented by its n × n adjacency matrix A, which consists of 0’s and 1’s. The (i,j)^th^ element of A is 1 if there is an edge connecting the i^th^ and the j^th^ node (e.g. if the correlation between the ith and jth regional volumes is greater than 0.5), and 0 otherwise. (The edges could be modeled as directed or undirected lines, but in this paper we focus only on undirected graphs). With the (i,j)th element of A as the actual correlation value if this value is greater than 0.5, and 0 otherwise, we get the adjaceny matrix of a weighted graph.

### Graph construction in the current analysis

Most of our analysis was done on binary graphs that modeled the statistical correlational relationship between regional volumes. There were two exceptions: the graph clustering and the computation of small world propensity (see below); in these cases we used weighted graphs. We will first describe the procedures that we adopted for binary graph generation, then we turn to the weighted graphs.

The first step in our analysis was to construct a binary graph by defining its nodes and edges. The nodes were the 87 brain areas included in the analysis, each subject providing one value of gray matter volume for every one of these areas. As described above, we removed the effects of age and gender by regressing these two factors on the structural volume of each region to obtain studentized residuals, which were then used for all further analyses. Next we needed to identify the edges. To do so, a simple procedure would be to calculate the absolute value of the Spearman’s correlation coefficient, rho (ρ), between the volume of pairs of areas and declare the nodes connected through an edge if the ρ value is higher than a set correlation threshold. However, no biological or clinical factor points towards an optimal threshold value for the correlation in a brain network. Second, one given individual correlation value corresponds to different *sparsity* levels (proportion of the number of edges out of total number of possible edges) in graphs obtained from patients and controls^14 19^ (Supplementary Fig. 1). On the other hand, only graphs with the same sparsity level can be meaningfully compared because sparsity influences graph properties. Therefore, we incremented sparsity in steps of 0.5%, from 0.5% to 99.5%. We obtained 199 sparsity levels, and for each of them we computed the 199 corresponding correlation thresholds for the patient and control groups, separately. Turning to the gray matter volumes that constituted our data set, we computed for each subject group a correlation between every pair of nodes. This correlation value was compared against the 199 correlation thresholds obtained as described above and an edge was declared in each of the 199 cases if the value of the correlation was equal or higher to the corresponding correlation threshold. For each subject group, we thus obtained 199 distinct binary graphs with 199 sets of parameters based on which we evaluated the AD-related alteration in brain topography. Because the sparsity levels ranged across the same values, this procedure ensured consistency in the graph comparisons across the patient-control groups. In particular, we would like to emphasize that in all the procedures described above the basis for edges in the graph construction was defined cross-sectionally, and not based on a time series as in fMRI data. We constructed binary graphs for control and patient groups in a similar manner after obtaining the sub-networks through the clustering algorithm.

Weighted graphs were utilized in two instances – first, when we partitioned the nodes based on a graph clustering algorithm and secondly, when we compared the small-world-propensity parameter between controls and patients. The reasons for using weighted graphs for these two instances are given in the corresponding sections (see below subsections *Graph clustering and Small World Propensity*). For the weighted graphs in our analyses, edges existed between all pair of nodes, and the weight for a particular edge was the absolute value of the Spearman’s correlation between the corresponding nodes.

### Global and sub-network analysis

We compared the properties of the graphs obtained from AD patients vs. normal controls in two different ways: globally, and after clustering the brain areas in two different sub-networks. For the global analysis, each AD patient (n = 100) or control subject (n = 135) provided 87 volumetric measurements for the corresponding nodes. For the second analysis, we repeated the computations after we first sorted the 87 brain regions in two distinct sub-networks. Our rationale was that in early AD, pathological processes may be more prevalent in some brain areas than in others, and focusing on more restricted networks may reveal important alterations in network properties that occur at this stage. We identified these sub-networks by applying a clustering procedure (described below) to a graph generated from the data belonging to 50 randomly selected subjects out of the 135 controls; the data from these 50 controls were then excluded from further analyses.

### Graph clustering

Characteristics of the structural brain network may differ between patients and controls on a local basis. Therefore, analysis of sub-networks may reveal important changes in patient’s brains that may otherwise escape identification when all regions are considered at once. To identify such local networks, graph clustering techniques can be used on data from the control subjects to delineate sub-networks of brain regions lowly connected among themselves but highly connected within. In our work, graph clustering was based on the divisive hierarchical clustering algorithm developed by Newman and Girvan^26^ which utilizes *edge betweenness* to group the nodes. We chose this clustering method because of thresholding issues. To cluster a binary graph we would have to first choose a threshold value (that is, one specific sparsity level). However, currently there is no biologically-based justification for using one sparsity level over the other. To avoid this problem, we applied the (widely used) partitioning technique for weighted graphs developed by Newman and Girvan to our analysis^26^.

The details of the algorithm are as follows. The betweenness score of an edge measures the number of shortest paths (paths between nodes with minimal number of vertices in between; see *Supplementary Text*) that include it. The intuitive idea behind the algorithm is that edges connecting separate clusters will have high edge betweenness scores because all the shortest paths from one cluster to another will have to traverse through them. Hence, deleting such edges will reveal the communities within the network. After deleting such edges in a first step, the edge betweeness scores are recalculated, and another deletion process occurs. Iteratively, the algorithm works as follows:

1. Calculate betweenness scores for all edges in the network.
2. Find the edge with the highest score and remove it from the network.
3. Recalculate betweenness for all remaining edges.
4. Repeat step 2.

The algorithm’s output is a dendrogram which represents an entire nested hierarchy of possible community divisions for the network. The point at which the dendrogram is cut is determined based on a modularity measure defined as the fraction of edges in the network that connect vertices of the same type minus the the expected value of the same quantity in a network with the same community divisions but random connections between the vertices. Values where this quantity is maximized indicate strong community structure, and it is at such values that the dendrogram is cut to obtain the graph clusters.

### Graph parameters

The graph parameters we used can be grouped under the following five large umbrellas: A) metrics of network efficiency; B) measures related to small worldness; C) measures of segregation; D) product measures of complexity; and E) entropy. Technically, measures in A) may be considered as small world properties or even measures of complexity; however, here we treated them separately because they have been conventionally considered as distinct measures. Mathematical descriptions of these parameters can be found in Supplementary Text.

*A. Metrics of network efficiency*, which measure the economical performance of the networks, were first defined in Latora and Marchiori,^33 34^ and further explored in brain network analysis by Achard and Bullmore^35^. Certain small world properties such as the average shortest path length can be calculated only for connected graphs, whereas the efficiency metrics can be calculated for disconnected graphs as well. Thus, these measures are especially useful at the low sparsity levels, where a graph could be disconnected.

- *Global efficiency* is a measure of how efficiently the structure of a network can support parallel information processing when all the nodes concurrently exchange packets of information^35^. Since there is strong prior evidence that brain supports massively parallel information processing, global efficiency is a biologically highly relevant measure for comparing patients and controls.
- *Cost efficiency*, the difference between global efficiency and sparsity level, is a measure that assesses the efficiency in relation to the sparsity of the network^35^. Since it has been shown previously that efficiency of a complex network increases as a function of its sparsity, it makes sense to account for sparsity, a measure of how inter-connected the nodes are, when assessing the effect of neurodegeneration in brain networks.

*B. Measures related to small worldness*. Small world networks are formally defined as networks in which the nodes are significantly more locally clustered, yet have approximately the same characteristic path length as random networks (networks where the nodes are connected at random)^36^. Each node of a small world network is highly likely to be connected to its nearest neighbors by a single edge (that is, high local clustering), but to get from one node to another node on the opposite side of the network, one need not traverse a large number of connections^15^. In other words, small-world networks are simultaneously highly segregated (compatible with modular/specific processing), yet highly integrated (compatible with distributed processing). Normal anatomical brain connectivity is thought to concurrently reconcile the opposing demands of functional integration and segregation, and hence is considered to have *small-world design*^37,15^.

- *Clustering Coefficient* measures the connectivity among adjacent nodes. For any node, it is calculated as the number of edges that exist between its nearest neighbors^38^. It is a measure of the extent of ‘cliquishness’ or the local clustering of the network^14^. For a brain network, this is a measure of segregated or modular processing.
- *Characteristic path length* (or *average shortest path length)* between any pair of nodes is the minimal number of edges that have to be traversed to reach from one node to the other. Averaged across all nodes, it is a measure of the extent of average connectivity or overall routing efficiency of the network^14^. For a brain network, this is a measure of integrated processing.
- *Sigma*, the ratio between the clustering coefficient and the shortest path length, is considered to be a summarized measure of small worldness. A network with small world property has large clustering coefficient and small characteristic path length, and hence the ratio, sigma will be large.
- *Small world propensity (SWP)* is a recently developed metric of small world property of a network^39^. The rationale for the new parameter was the heavy dependence of both characteristic path length and clustering coefficient on the sparsity level of the network. The main advantage of SWP over *Sigma* is that it is relatively independent of the sparsity level of the network and thus it is a more robust measure of small world property. Roughly speaking, SWP indicates how much the clustering coefficient and characteristic path length of a given network differ from the corresponding values in both lattice and random networks, constructed with the same number of nodes and the same degree distribution. SWP ranges between 0 and 1, with larger values indicating more small-world-ness; networks with values below a reference value of 0.6 were considered to be exhibiting only weak evidence of small world property. Although a binary version and a weighted version of SWP were presented in Muldoon et al. (2015), we used the weighted SWP in our analysis, since the main rationale behind SWP is to avoid dependence on sparsity levels.

*C. Measures of segregation*

- *Betweenness centrality* is defined as the number of shortest paths that run through a given node^40^. Averaged across all nodes, it is a measure that captures the influence of a node over information flow between other nodes in the network.

*D. Product measures of complexity* are based on the idea that networks with intricate structural patterns of connections have a medium number of edges; that is, they are neither very sparse nor highly connected. It has been established that both minimally connected networks and fully connected networks have complexity values approximately zero^41^. The two product measures described below are products of two factors: F_1_ × F_2_. F_1_ assigns values near zero for sparsely connected networks and large values for highly connected networks. In contrast, F_2_ assigns values near zero for highly connected networks and large values for sparse networks. Therefore, the product F_1_ × F_2_ will always have small values for both sparse and highly connected networks and large values for networks with medium number of edges. In the context of this analysis, a smaller value for either of these two parameters corresponds to a decrease in the complexity of connectivity between brain areas.

- *Medium Articulation for graphs* is the product of two factors: Redundancy, R and mutual information I. R is zero for the sparse graph, but maximum for the fully connected graph; I varies in the opposite way^41^.
- *Graph Index Complexity* is proportional to the product between the normalized index, c_r_, of the largest eigenvalue of the adjacency matrix of a graph, and its difference from unity, 1-c_r_. c_r_ values ranges from 0 to 1; it is 0 for a minimally connected graph and 1 for maximally connected graph, so that the product measure c_r_(1-c_r_) is largest for graphs with medium number of edges, a characteristic of a complex graph^41^.

*E. Entropy* in thermodynamics is defined as the amount of disorder; that is, the number of specific ways in which a thermodynamic system may be arranged. In a similar vein, the entropy of a complex network quantifies its structural diversity and is referred to as the information content of a graph. The entropy parameters considered in this paper are described below.

- *Topological Information Content (TIC):* The information content of a network largely depends on the arrangement of the nodes and edges of the network. The TIC measure considered in our analysis is small for graphs in which each node has the same number of neighbors, and it attains maximum for asymmetric graphs^42 43^.
- *Bertz Index (BI):* Reasoning that complexity should increase with the number of nodes regardless of whether they are all equivalent, Bertz introduced a measure of entropy as a function of the number of nodes and edges^44^.
- *Vertex Degree Information-Equality Based Information Index* is an entropy measure calculated based on partitioning the nodes of a network based on their degrees (number of adjacent nodes). A graph with an approximately uniform degree distribution will have smaller values for this index compared to graphs with non-uniform degree distributions^45 46^.
- *Graph Vertex Complexity Index* measures the average heterogeneity of the *distances* associated with each node in a network. A *distance* is a quantification of the number of edges that link a pair of nodes. For example, adjacent nodes have a distance of 1. Higher values of this parameter indicate a more complex network^47^.
- *Mean Information content on the Edge Equality* quantifies the heterogeneity of all the distances in a network. This parameter is thus very similar to the graph vertex complexity index, except that rather than looking at individual nodes, it considers the network as a whole^46^.
- *Mean Information Content on the Edge Magnitude* quantifies the *heterogeneity of the magnitudes* of these distances. Higher values of this parameter indicate a more complex network^46^.
- *Off Diagonal Complexity*: It has been suggested that a complex graph has many different entries in its so-called node-node link correlation matrix *c_ij_*, where *c_ij_* denotes the number of all neighbors with degree j ≥ I of all nodes with degree i. Off diagonal complexity measures this diversity. More specifically, off diagonal complexity is high for a graph where the nodes of a given degree have no preference for the degree of their neighbors^41^.

### Statistical Analysis

Comparison between groups were performed using a t-test as well as a permutation test. By varying the sparsity levels (see above), we obtained a range of values for each graph parameter. This procedure was run separately for controls and patient groups, resulting in the same number of graph parameter values for each of the two cases. The two data sets were compared by using a simple t-test. We also used a permutation test with 10,000 iterations, as suggested by Bassett and collaborators^48^. For each graph parameter, with the range of values for each parameter obtained for various sparsity levels on the y-axis, and the corresponding sparsity level on the x-axis, one obtains two curves, one for the patient group and another for the control group. The absolute difference between the two curves (that is, the area between the two curves) was the statistic used in the permutation test. In order to conduct the permutation test, at each iteration, the labels indicating patients and controls were randomly permuted, and the area between the curves was calculated for two pseudo-groups of ‘patients’ and ‘controls’. P-value for the permutation test was the proportion among these 10,000 area values that was greater than the area value obtained for the original patients and controls. Adjustments for multiple comparisons used Benjamini-Hochberg’s procedure to control the False Discovery Rate (FDR)^49^. We considered a parameter to be significantly different between patients and controls only if the FDR-adjusted p-values for both the t-test and the permutation test were significant at a 0.05 level.

## RESULTS

### Overall network analysis

The graphs of the overall networks were similar in AD patients and control groups, and the analysis did not reveal significant differences between any of their parameters (Fig. 1A, Table 2). Thus, this analysis suggested that in the early stages of AD, the graph properties of large structural networks do not change even as degenerative processes may already have caused significant neural damage. However, since neurodegeneration occurs at different rates in different areas, an overall analysis may miss modifications taking place in more localized networks. Our subsequent graph analysis, applied after clustering the brain areas, was in agreement with this idea.

**Figure 1.**
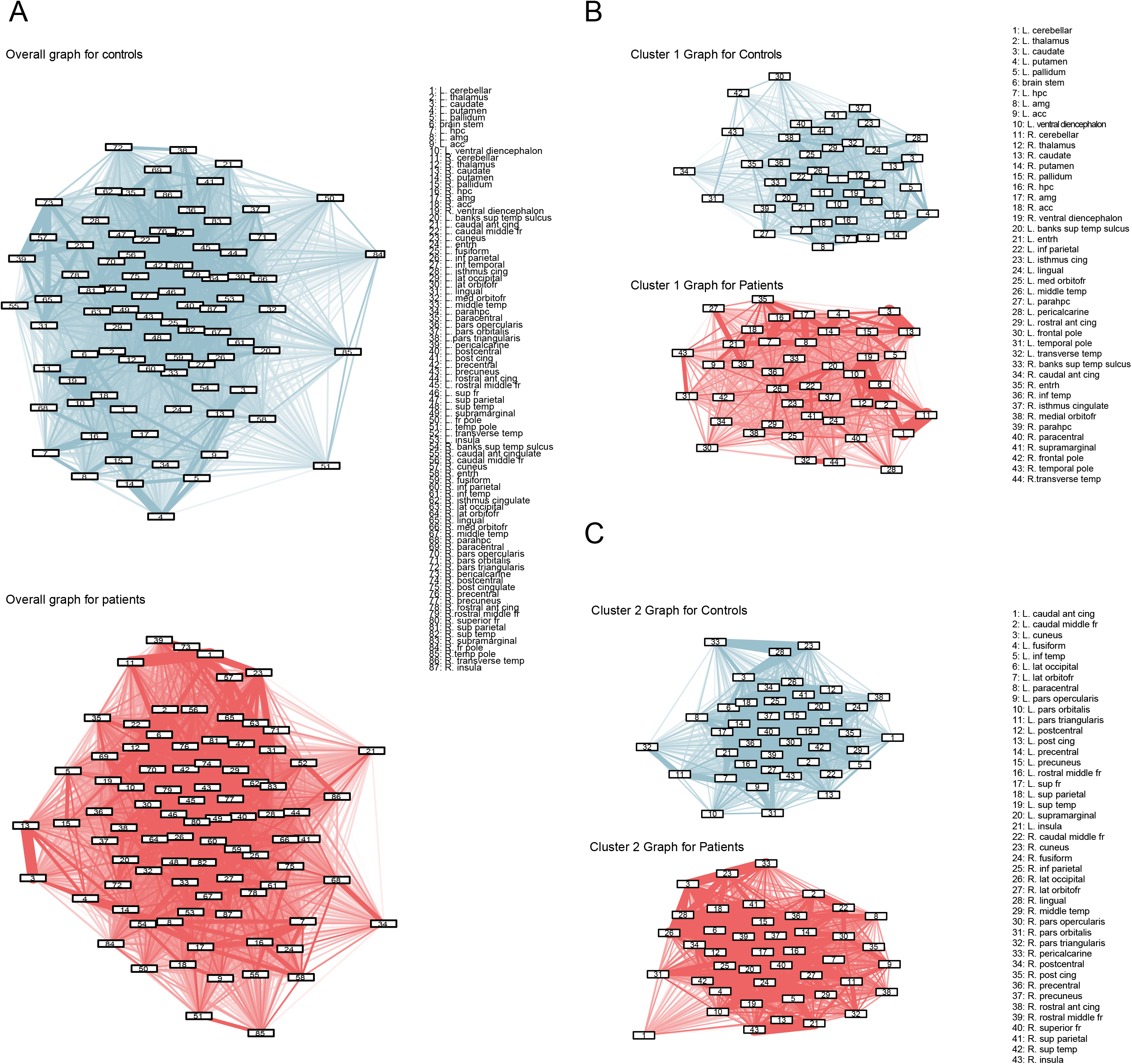
**Graphs constructed from volumes of gray matter data.** Graphs in blue represent the networks for controls and graphs in red represent networks for patients. Sparsity level for all graphs is 50%. **A. Graphs of all brain areas.** These graphs include all 87 regions included in this analysis. The properties of the two graphs were similar. **<u>B and C</u>. Graphs after grouping the brain areas in two sub-networks.** Each cluster contained areas whose volume diminished, but hippocampus and entorhinal cortex, areas known to be markedly affected even in early stages of Alzheimer’s, were included in the first sub-network. The two sets of graphs showed significant differences in their properties when comparing control and AD patient groups; and the pattern of the changes was different in sub-network 1 (B) vs. sub-network 2 (C).

**Table 2.** **Results of the graph analysis applied to all brain areas considered as a single large network.** Data are reported as mean and (standard deviations). AD = Alzheimer’s

|  | Parameter | Overall |  |  |  |
| --- | --- | --- | --- | --- | --- |
|  |  | Controls<br>Mean<br>(stdev) | AD's<br>Mean<br>(stdev) | p-value(s)<br>1) t-test<br>2) permutation<br>based | p-value<br>adjusted<br>for FDR<br>1) t-test<br>2)<br>permutati<br>on based |
| Metrics of<br>network<br>efficiency | Global Efficiency | 0.640<br>(0.321) | 0.643<br>(0.317) | 1) 0.9392<br>2) 0.5060 | 1) 0.9625<br>2) 0.8385 |
|  | Cost Efficiency | 0.150<br>(0.518) | 0.153<br>(0.517) | 1) 0.9625<br>2) 0.5060 | 1) 0.9625<br>2) 0.8385 |
| Measures<br>related to<br>small<br>worldness | Clustering coefficient | 0.713 (0.157) | 0.695<br>(0.159) | 1) 0.2602<br>2) 0.3468 | 1) 0.9625<br>2) 0.8385 |
|  | Average shortest pathlength | 1.709 (0.594) | 1.716<br>(0.634) | 1) 0.9165<br>2) 0.8675 | 1) 0.9625<br>2) 0.9295 |
|  | Sigma | 0.487 (0.226) | 0.476<br>(0.226) | 1) 0.6471<br>2) 0.5119 | 1) 0.9625<br>2) 0.8385 |
| Measure of<br>segregation | Betweenness centrality | 25.53 (18.98) | 25.77<br>(19.08) | 1) 0.9033<br>2) 0.9655 | 1) 0.9625<br>2) 0.9655 |
| Product<br>Measures | Medium Articulation | 0.333 (0.317) | 0.342<br>(0.321) | 1) 0.7773<br>2) 0.4682 | 1) 0.9625<br>2) 0.8385 |
|  | Graph Index Complexity | 0.713 (0.279) | 0.718<br>(0.272) | 1) 0.8498<br>2) 0.5535 | 1) 0.9625<br>2) 0.8385 |
| Entropy | Topological Info Content | 6.196 (0.628) | 6.192<br>(0.638) | 1) 0.9544<br>2) 0.7169 | 1) 0.9625<br>2) 0.8670 |
|  | Bertz Index | 1099.6 (54.6) | 1099.3<br>(55.5) | 1) 0.9544<br>2) 0.7169 | 1) 0.9625<br>2) 0.8670 |
|  | Vertex Degree Information-Equality Based Information<br>Index | 4.808 (0.801) | 4.833<br>(0.775) | 1) 0.7635<br>2) 0.7514 | 1) 0.9625<br>2) 0.8670 |
|  | Graph Vertex Complexity<br>Index | 1.099 (0.508) | 1.056<br>(0.456) | 1) 0.3901<br>2) 0.5517 | 1) 0.9625<br>2) 0.8385 |
|  | Mean Information content<br>on the Edge Equality | 8.000 (1.360) | 8.123<br>(1.328) | 1) 0.3663<br>2) 0.5590 | 1) 0.9625<br>2) 0.8385 |
|  | Mean Information content<br>on the Edge magnitude | 10.36 (1.31) | 10.40<br>(1.31) | 1) 0.7666<br>2) 0.2867 | 1) 0.9625<br>2) 0.8385 |
|  | Off Diagonal Complexity | 0.738 (0.142) | 0.754<br>(0.144) | 1) 0.2747<br>2) 0.3394 | 1) 0.9625<br>2) 0.8385 |
| New sensitive measure for small world properties based on weighted graph |  |  |  |  |  |
|  | SWP | 0.1154<br>(0.0943) | 0.1918<br>(0.1081) | 0.5211 |  |

### Sub-network analysis

#### Clusters

We first identified the brain sub-networks based on data from 50 randomly chosen normal subjects. As clustering algorithm, we used a divisive algorithm that estimated the number of clusters in our dataset as being equal to two^26^. The two clusters were constituted of 44 and 43 regions, respectively (Table 1, Figs. 1B, and 2). Each sub-network included areas whose gray matter volumes decreased in AD patient group (Fig. 3, Table 3), but notably, areas of the medial temporal lobe known to be among the most affected in early stages of AD (hippocampus, amygdala, entorhinal cortex, parahippocampal cortex) were grouped in the first sub-network.

**Figure 2.**
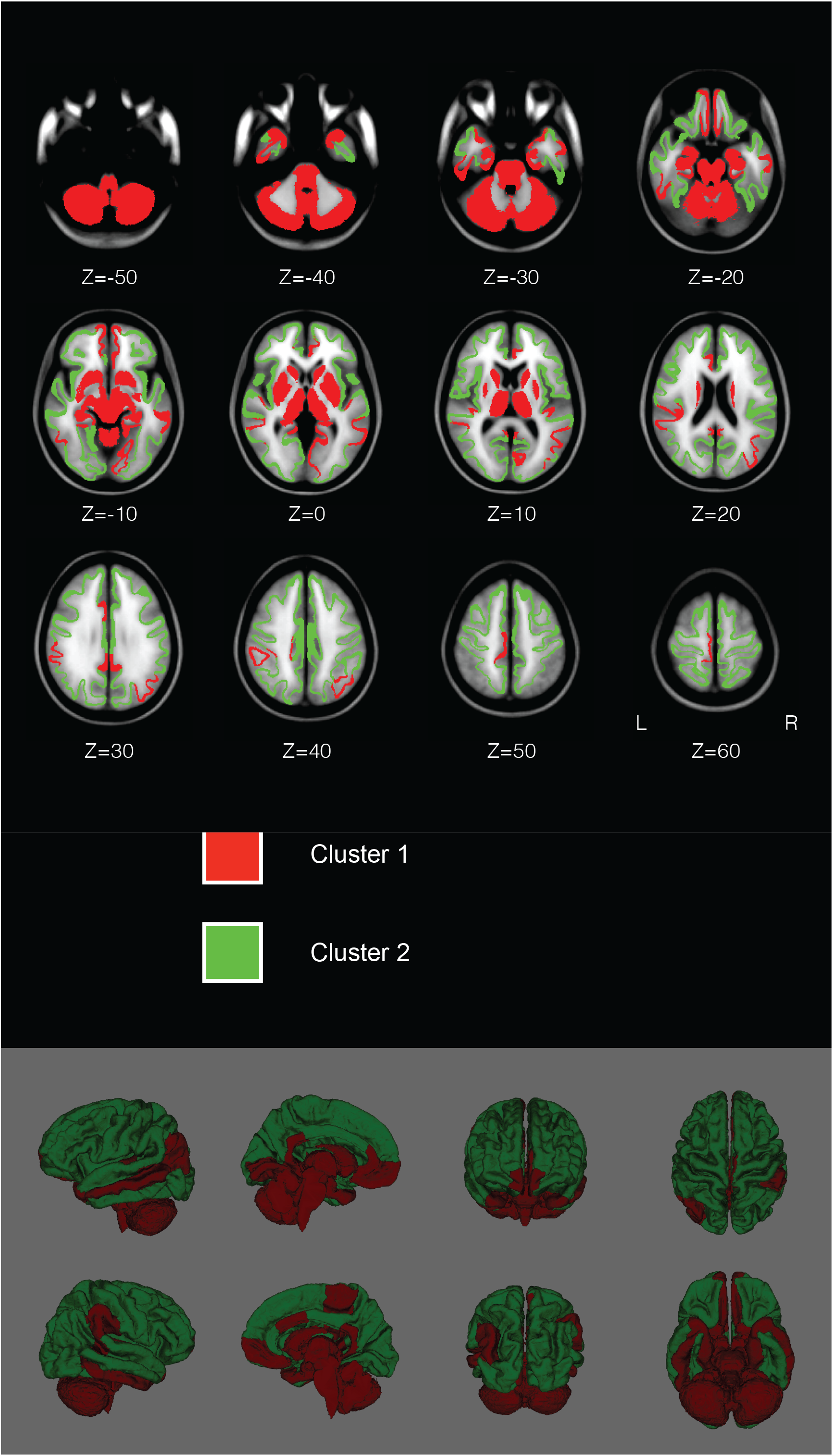
**FreeSurfer 5.3.0 based brain mapping of the structural regions’ clusters.** The two clusters of brain areas and their hubs are shown in distinct colors on successive horizontal sections (Z = z axis) from the ventral (top left) to the dorsal (bottom right) through a human brain. Starting from the left, the bottom of the figure shows the areas of the two clusters from lateral (left-right), medial (left-right), anterior-posterior, and dorsal-ventral views of the brain, respectively. The network properties of these clusters were significantly different in AD patients when compared to healthy elders.

**Figure 3.**
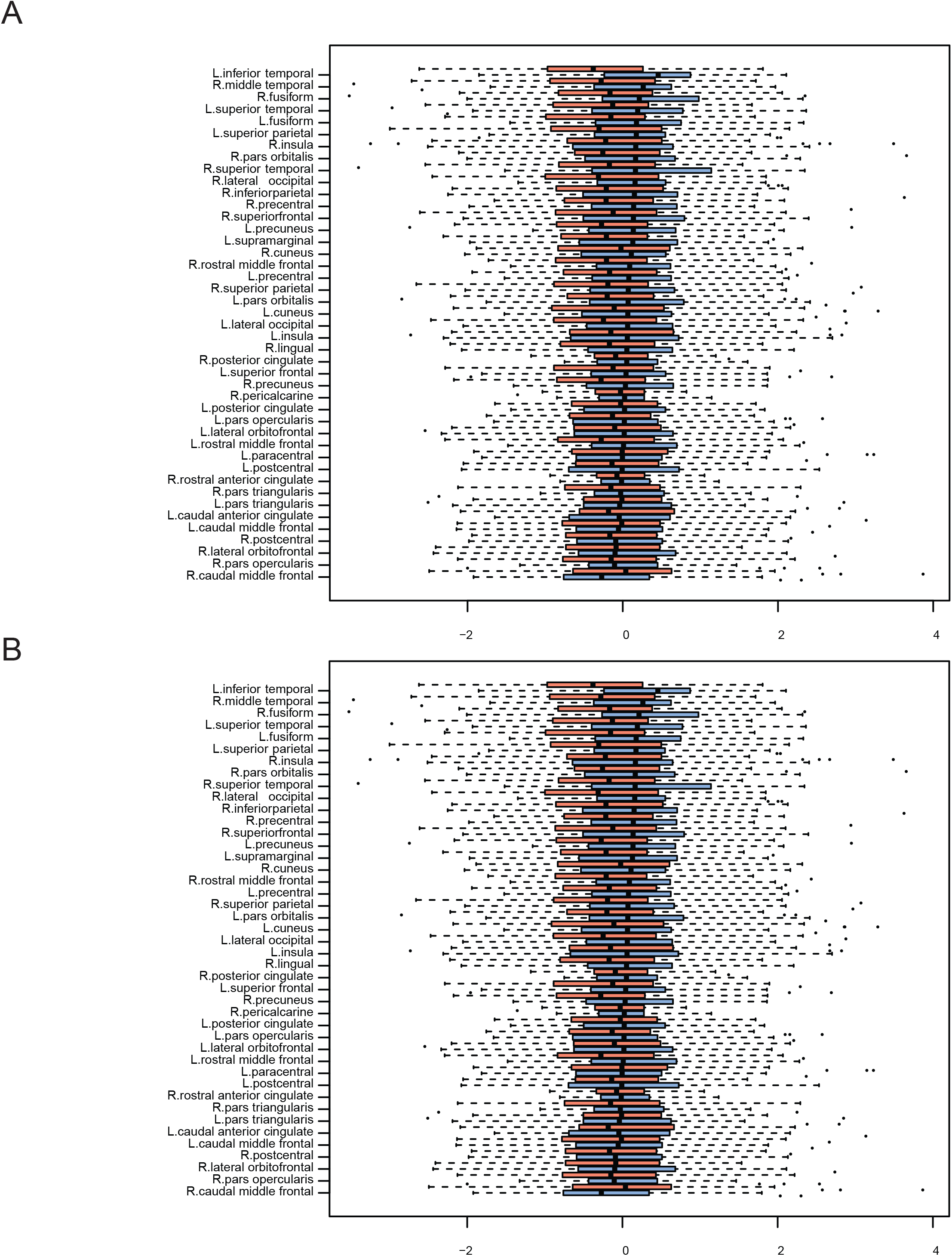
**Boxplots of gray matter volumes in control (blue) and patient (red) groups for the two clusters.** A = Sub-network1; B = Sub-network 2. For each cluster, data are organized based on the values of the control group sorted in descending order from top to bottom. Data from the AD patients are shown in a box plot located immediately above the corresponding control group. To reveal the volumetric changes in brain areas, dark vertical lines indicate the medians of the brain regions for the control and patient groups, respectively. Values located to the left of the zero mark indicate a smaller volume. The plots reveal that both sub-networks contained brain areas affected by neurodegeneration.

**Table 3.**
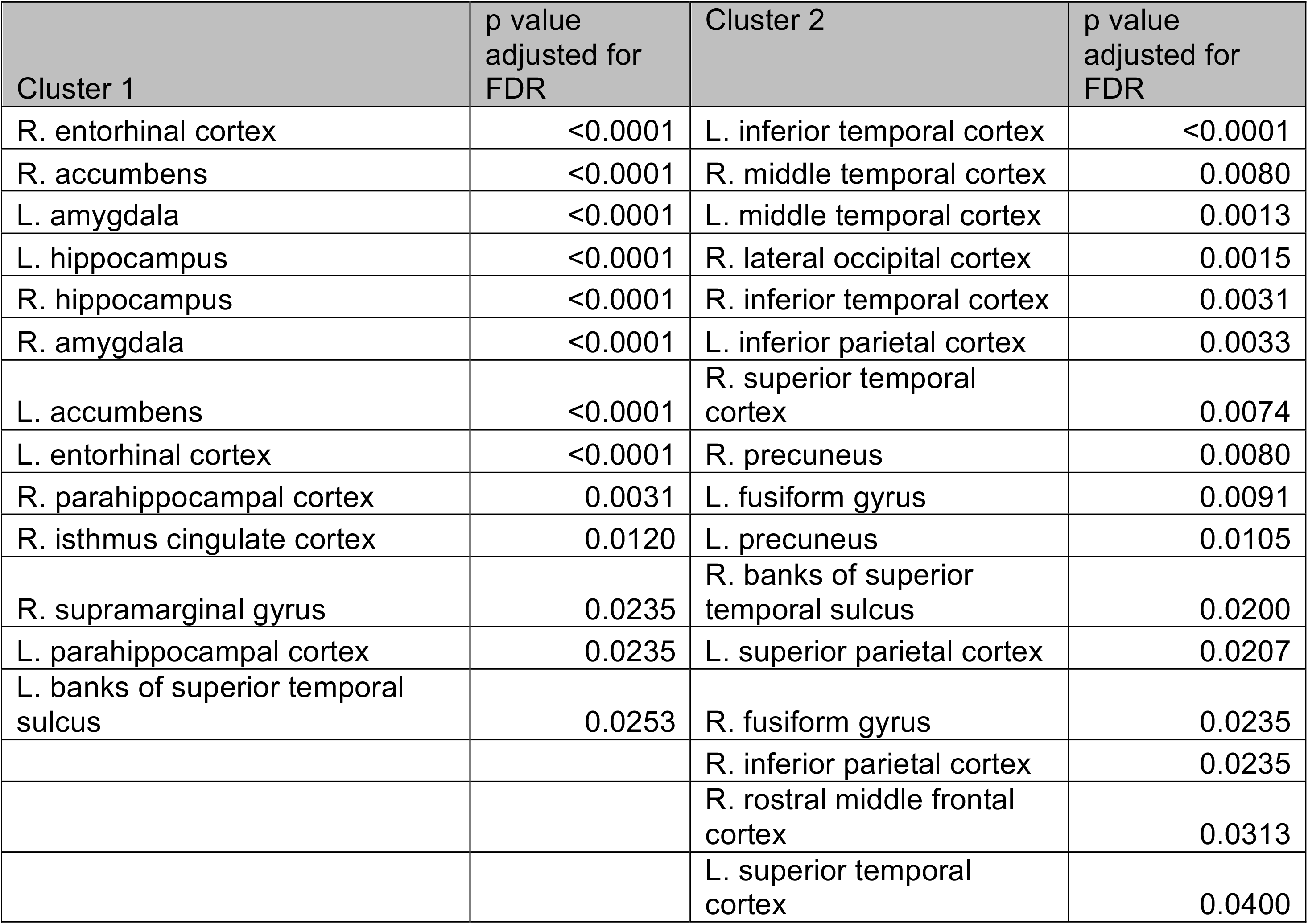
**Areas with significant shrinkage of gray matter volume.** Raw *p* values obtained from the Bayesian analysis as well as the two-sample comparison of the residual volume of regions between the two clusters were adjusted for multiple testing using the ‘multtest’ package in R. Because Bayesian analysis involved comparing 15 graph parameters between patients and controls, and latter analyses involved comparing 87 regions between the two clusters, multiple testing adjustments were done using the Benjamini-Hochberg (BH) method controlling for the false discovery rate (FDR).

### Changes in the sub-networks

The results we obtained in the analysis of the large brain network may be due to the fact that at the early AD stage, neurodegenerative processes do not have a sufficiently marked effect to introduce modifications in network topography, but this was unlikely because shrinkage of gray matter volumes was quite prominent in our data set (Fig. 3, Table 3). Alternatively, neurodegeneration may affect the gray matter volumes, but preserve the topography of the network. This hypothesis is equally unlikely because particularly in early AD, different brain areas are altered heterogeneously – some more, such as in the medial temporal cortex, and some less, such as in the prefrontal areas. The more likely scenario is that assembling all areas in one large network may mask real and important changes in the structure of more circumscribed brain networks. Supporting this idea, a different pattern of results emerged when we applied the graph analysis to the two sub-networks identified through the clustering procedure. We compared the mean value and the entire curve of each graph parameter (obtained across sparsity levels), after we adjusted the p-values to correct for multiple comparisons. The means, standard deviations, p-values, and adjusted p-values are presented in Table 4. We considered a significant change in the parameter only if both the mean and the curve comparison indicated statistical significance after adjustment for multiple comparisons. Fig. 4 graphically illustrates the variation of these parameters between control and patient groups in the two sub-networks with statistically significant differences highlighted in gray. As opposed to the results of the large network analysis, which did not suggest changes, in this case we found that compared to control condition, AD was associated with alterations in each of the two sub-networks, and surprisingly, there was in fact some indication that the two sub-networks were affected differently. The properties of the sub-network that included medial temporal lobe areas were modified in a direction consistent with a deterioration of network small-world properties and loss of network diversity and complexity in AD patients. In contrast, changes in the second sub-network, while also reflecting a decrease network diversity and complexity, revealed modifications suggesting an increase in small world properties.

**Table 4.** **Results of the graph analysis applied to the two sub-networks.** Data are reported as mean and (standard deviations). AD = Alzheimer’s Disease *, **, and *** = statistically significant

|  | Parameter | Cluster 1 |  |  |  | Cluster 2 |  |  |  |
| --- | --- | --- | --- | --- | --- | --- | --- | --- | --- |
|  |  | Controls<br>Mean<br>(stdev) | AD's<br>Mean<br>(stdev) | p-value(s)<br>1) t-test<br>2) permutation based | p-value adjusted for FDR<br>1) t-test<br>2) permutation based | Controls<br>Mean<br>(stdev) | AD's<br>Mean<br>(stdev) | p-value(s)<br>1) t-test<br>2) permutation based | p-value adjusted for FDR<br>1) t-test<br>2) permutation based |
| Metrics of network efficiency | Global Efficiency | 0.6362<br>(0.3951) | 0.5820<br>(0.4913) | 0.2306<br><0.0001 | 0.2705<br><0.0001* | 0.5022<br>(0.3292) | 0.4600<br>(0.3481) | 0.2187<br>0.2420 | 0.6756<br>0.3633 |
|  | Cost Efficiency | 0.1462<br>(0.4689) | 0.0920<br>(0.4773) | 0.2585<br><0.0001 | 0.2705<br><0.0001* | 0.0123<br>(0.5800) | -0.0300<br>(0.5899) | 0.4759<br>0.2422 | 0.7915<br>0.3633 |
| Measures related to small worldness | Clustering coefficient | 0.6519<br>(0.1644) | 0.5742<br>(0.1725) | <0.0001<br><0.0001 | **0.0001\*\*\***<br>**<0.0001\*\*\*** | 0.8197<br>(0.1335) | 0.8081<br>(0.1651) | 0.4472<br>0.7719 | 0.7915<br>0.8906 |
|  | Average shortest pathlength | 1.7864<br>(0.6180) | 1.8703<br>(0.6155) | 0.1797<br><0.0001 | 0.2572<br><0.0001*** | 1.3890<br>(0.3905) | 1.3808<br>(0.4403) | 0.8464<br>0.9845 | 0.9069<br>0.9845 |
|  | Sigma | 0.4259<br>(0.2116) | 0.3628<br>(0.2088) | 0.0032<br><0.0001 | **0.0161\***<br>**<0.0001\*\*\*** | 0.6478<br>(0.2191) | 0.6538<br>(0.2449) | 0.7998<br>0.4047 | 0.9069<br>0.5519 |
| Measure of segregation | Betweenness centrality | 13.51<br>(8.71) | 14.60<br>(10.80) | 0.2702<br>0.0047 | 0.2705<br>0.0055** | 7.6067<br>(7.0972) | 6.8102<br>(6.5110) | 0.2489<br>0.6451 | 0.6756<br>0.8064 |
| Product Measures | Medium Articulation | 0.4421<br>(0.3432) | 0.4807<br>(0.3490) | 0.2705<br><0.0001 | 0.2705<br><0.0001* | 0.1985<br>(0.2684) | 0.1998<br>(0.2807) | 0.9622<br>0.1504 | 0.9622<br>0.3020 |
|  | Graph Index Complexity | 0.7169<br>(0.2488) | 0.6609<br>(0.2782) | 0.0366<br><0.0001 | 0.1097<br><0.0001* | 0.5538<br>(0.3310) | 0.5056<br>(0.3571) | 0.1678<br>0.0082 | 0.6756<br>0.0409* |
| Entropy | Topological Info Content | 5.1788<br>(0.6390) | 5.0828<br>(0.7933) | 0.1886<br>0.0036 | 0.2572<br>0.0044** | 4.8453<br>(0.9217) | 4.4341<br>(1.2324) | 0.0002<br>0.0033 | **0.0017\*\***<br>**0.0245\*** |
|  | Bertz Index | 468.08<br>(28.12) | 463.86<br>(34.90) | 0.1886<br>0.0485 | 0.2572<br>0.0485* | 441.67<br>(39.63) | 423.99<br>(52.99) | 0.0002<br><0.0001 | **0.0017\*\***<br>**<0.0001\*\*\*** |
|  | Vertex Degree<br>Information-Equality Based<br>Information<br>Index | 3.8720<br>(0.6533) | 3.6288<br>(0.7486) | 0.0007<br><0.0001 | **0.0052\*\***<br>**<0.0001\*\*\*** | 3.5937<br>(0.7643) | 3.4940<br>(1.0042) | 0.2702<br>0.0164 | 0.6756<br>0.0540 |
|  | Graph Vertex<br>Complexity<br>Index | 1.2584<br>(0.4964) | 1.1921<br>(0.4745) | 0.1782<br>0.0083 | 0.2572<br>0.0089* | 0.8761<br>(0.4322) | 0.8474<br>(0.4651) | 0.5276<br>0.8512 | 0.7915<br>0.9120 |
|  | Mean<br>Information<br>content on the<br>Edge Equality | 6.1804<br>(1.1662) | 5.8805<br>(1.4035) | 0.0223<br><0.0001 | 0.0835<br><0.0001*** | 5.7869<br>(1.1879) | 5.6901<br>(1.6588) | 0.5081<br>0.0180 | 0.7915<br>0.0540 |
|  | Mean<br>Information<br>content on the<br>Edge<br>magnitude | 8.3339<br>(1.1572) | 8.1127<br>(1.2331) | 0.0685<br><0.0001 | 0.1467<br><0.0001* | 9.1248<br>(0.8405) | 9.0940<br>(0.9515) | 0.7351<br>0.1611 | 0.9069<br>0.3020 |
|  | Off Diagonal<br>Complexity | 0.6480<br>(0.1594) | 0.6140<br>(0.1977) | 0.0621<br>0.0036 | 0.1467<br>0.0044** | 0.6535<br>(0.1339) | 0.6456<br>(0.1896) | 0.6339<br>0.1488 | 0.8644<br>0.3020 |
| New sensitive measure for small world properties based on weighted graph |  |  |  |  |  |  |  |  |  |
|  | SWP | 0.1983<br>(0.0844) | 0.1740<br>(0.0780) | 0.8882 |  | 0.1288<br>(0.0455) | 0.2588<br>(0.0341) | 0.0511* |  |

**Figure 4.**
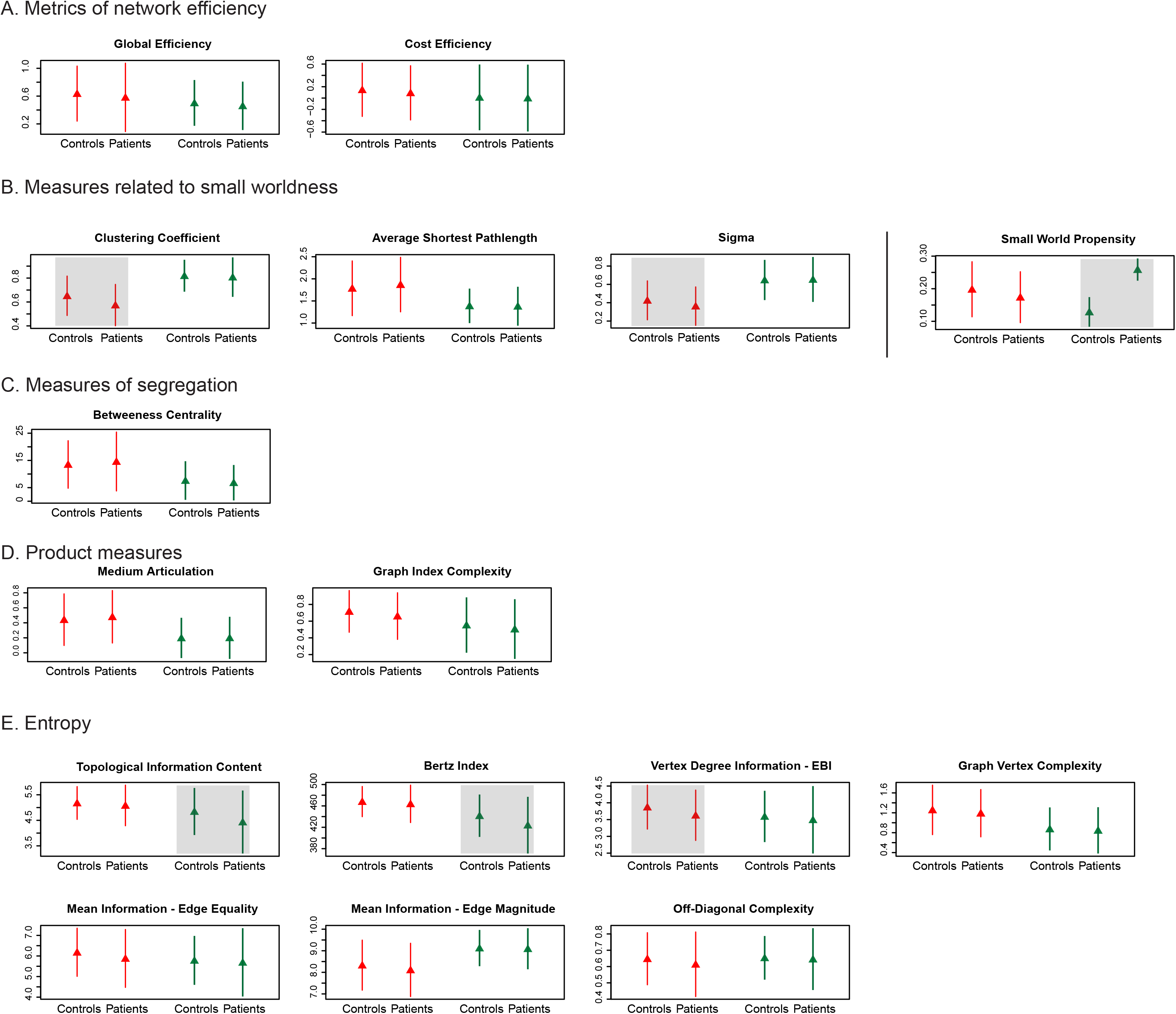
**Changes in graph properties for the two sub-networks.** The data shown in each plot correspond to values presented in Table 4. The gray highlights mark effect sizes with associated probabilities smaller than 0.5 in both t-test and permutation test. Overall, in the first sub-network (red), which included the hippocampus and ento-rhinal cortex, the variation in parameter values indicated deterioration of small world characteristics and diminished complexity and diversity in the patient group when compared to control. In the second sub-network (blue), small world properties were not affected negatively but rather there was an improvement as measured by SWP. As in the first cluster, complexity and diversity also diminished.

In the first sub-network there was no significant change in efficiency measures with AD (Fig. 4A). In contrast, both the clustering coefficient and sigma declined with AD while the characteristic path length did not change significantly (Fig. 4B). Sigma, considered a summary of small world properties of a network, is calculated as the ratio between the clustering coefficient and the characteristic path length. Therefore, these results suggested that AD altered the normal balance between segregation and integration of information processing by diminishing the former (a decrease of connectivity among adjacent nodes is a sign of smaller node clustering and hence ability to segregate information processing). The nodes’ ability to modulate information flow (betweenness centrality, Fig. 4C) did not change, and neither did the complexity of connectivity between brain areas (product measures, Fig. 4D). There was however an indication that the structural diversity of the sub-network decreased (significant diminishing of vertex degree information, Fig. 4E). Overall, the results suggested that in the first sub-network, AD-related changes uniformly pointed towards deterioration of network properties.

In the second sub-network, AD seemed to also alter the network properties, but in this case the pattern was different. As in the first sub-network, there were no changes in network efficiency, segregation or product measures of complexity (Fig. 4A, C, D), and the diversity of the network decreased with AD (lower topological information content and Bertz index, Fig. 4E). However, the small world property of this sub-network seemed to be altered differently: while clustering coefficient, average pathlength, or sigma did not vary significantly, the SWP parameter suggested an *increase* in small world properties (Fig. 4, B). Overall the results suggested that the AD-induced alterations had multifaceted consequences, and that perhaps, at these incipient stages, deterioration of some brain areas triggers modifications aimed at recovering normal functional balance in some other brain areas.

## DISCUSSION

We applied complex network analysis to investigate the structural modifications that accompany early AD in 87 brain volumes obtained from FreeSurfer automatic segmentation. When all the brain areas under consideration were used to form one large network, the analysis did not identify differences in the network properties between control and AD patients. However, because in early AD neurodegenerative processes affect individual brain areas to different degrees^50^, we also focused our investigation on more restricted networks by partitioning the 87 brain regions into two sub-networks. In contrast to the results of the global network analysis which did not signal any changes, the graph analysis applied to these sub-networks revealed marked differences between control and AD patient groups. In the first sub-network, which encompassed medial temporal lobe areas such as hippocampus, entorhinal cortex, and parahippocampal cortex, data from AD patients showed small world parameters consistent with a reduced ability to concomitantly engage in segregated and integrated information processing. Extending the analysis to entropy, we additionally found evidence of reduced structural diversity (smaller vertex degree information-equality based entropy in AD group). The second sub-network also showed deterioration of complexity/diversity, although for different entropy parameters (TCI and Bertz index). Because TCI, Bertz index, and vertex degree information-equality based index are all closely related, and because differences and similarities among entropy parameters are not clear^51^, the most likely interpretation of these results is that in both sub-networks, AD effects are similar, namely a decrease in complexity and diversity. This conclusion is supported by the trend in the rest of the parameters of the entropy group, although these changes were not statistically significant. In the second sub-network however, we found no AD-related reduction in small world properties. In fact, a recently developed parameter, SWP, suggested an *increase* in small world properties with AD, revealing opposing trends in the two sub-networks. Our analysis therefore suggests that in early AD, neurodegenerative processes modify the topology of brain networks in complex ways.

### Neurodegeneration has heterogenous effects on network properties

Both sub-networks considered for this analysis included areas that showed diminished gray matter volume in AD, and yet the properties of these sub-networks were not affected homogenously: the complexity and diversity of the two sub-networks decreased, but the effects on the small world properties seemed to evolve in opposite directions. To the best of our knowledge, this is the first study associating AD with structural connectivity modifications that alter the diversity and complexity within brain networks. Second, our findings indicate that one single process – neurodegeneration – can have distinct effects in different networks. Here, we cannot establish whether these effects were due to the intrinsic topology of each of the two sub-networks, to specificities of neurodegeneration in the individual areas that constituted these two sub-networks, or to a combination of these two possibilities. What our results highlight is that encompassing multiple measures to quantify changes in brain networks reveals important information regarding the nature of the alterations. Many of the large, sparse biological networks found in nature, including brain networks, are small world networks^36^, whose characteristic is an optimal balance between local specialization (specialized processing in distinct regions) and global integration (rapid combination of processing across distributed regions). Small world networks achieve this equilibrium by combining high clustering of the nodes with short path lengths between them. Evaluating the small world properties of a network can thus provide important information about the functionality of that network. However, small world properties provide only one window into the properties of the intricate entities that brain networks are. Other important aspects of a network are its diversity, quantified by *entropy* (the higher the entropy, the higher the diversity) and its *complexity*. Both these parameters rank high in functional large structural networks. The results of our analysis suggested that in early AD the ubiquitous deterioration of complexity/diversity of the networks is nonetheless coupled with a heterogenous effect on the small world properties. Neural circuits are plastic, and thus in early AD, modifications in the areas first affected may trigger reorganization processes in other areas, possibly the ones with still intact structure and function. Do empirical data support this hypothesis? If so, what is the nature of the reorganization processes? Does this remain true in advanced AD, where neurodegeneration is profound and encompasses vast portions of the brain? Further investigations extended to include multiple parameters and patients with stages of AD ranging from mild to advanced would provide valuable insights into the dynamics of this disorder.

### Brain areas included in network construction influence assessment of modifications in network properties

Our results show that in early AD, modifications in the properties of brain networks are prominent in delimited sub-networks. We grouped the gray matter volumes of the 87 brain regions of our data set by using a graph clustering method based on the divisive hierarchical clustering algorithm with modularity maximization. The advantage of this clustering procedure is that it can be applied to weighted graphs. Other approaches for generating the sub-networks could also be used, for example ones that focus on sub-networks of areas *a priori* known to be anatomically or functionally interconnected. The point we want to emphasize here is that investigating the properties of a structural network constructed from a large number of brain areas may not reveal important alterations in properties of more local sub-networks. This may be particularly important for early AD, when degenerative processes are not very advanced and have not uniformly affected all brain regions. The general conclusion is that when applying graph analysis, the brain areas included in graph construction is a factor that should be taken into account in evaluating the results.

### Early AD is associated with modifications in measures related to small world properties

In our analysis, both sub-networks showed significant alterations in parameters related to small world properties. We found that AD processes modified the first sub-network so as to diminish their small world properties, consistent with other previous studies reporting altered small world properties^12,13^ (but see^52^). However, the analysis of the second sub-network suggested that the modifications in the patient group went in quite the opposite direction: AD was associated with an increase in the SWP parameter, considered to be a sensitive indicator of small world properties^39^. This parameter also suggested a non-significant trend towards reduction of small world properties with AD in the first sub-network. Thus, at least in early AD, aside of the negative effects of the neurodegeneration on small world properties of brain networks, there may be some other changes taking place concomitantly which do not necessarily result in negative effects on the network parameters. While our analysis indicates that AD indeed reduces the small world properties of at least some structural brain networks, understanding the precise processes causing this phenomenon requires more extensive investigations.

### Functional implications of network modifications in AD

Our analysis was not focused on the default mode network (DMN), a collection of brain areas whose function has been linked to performance in tasks that require cognitive demand^53^ and whose activity is reduced in AD^54^. The functional role of DMN is presently not fully understood, but its constituent areas are known to be involved in episodic memory, which is the type of memory that declines early in AD. Because DMN’s functional connectivity is linked to the network’s structural connectivity^55^ and because AD seems to spread along brain networks^56 57 52^, structural modifications of DMN in AD are likely to have functional implications. In our analysis, DMN areas (medial temporal lobe, posterior cingulate cortex, medial prefrontal cortex, inferior parietal cortex) were spread across the two sub-networks we identified. One possibility that can be further investigated is suggested by the fact that hippocampus, amygdala, and parahippocampal cortex are found in the first sub-network, while inferior parietal cortex and right medial orbitofrontal cortex are found in the second sub-network. One may speculate that as the medial temporal lobe areas deteriorate in the early stages of AD, changes also take place concomitantly in the rest of the areas, which are affected at later points of the disease^57 50,58,59^. Further studies of the dynamics of modifications taking place during the long course of AD would promote our understanding of pathological processes in AD and inform the design of approaches in AD treatment.

## CONCLUSION

In this study, we quantitatively analyzed the small-world properties, efficiency, complexity and diversity of structural brain networks in Alzheimer’s patients and healthy elders using data provided by structural MRI. Our results indicated that AD-related modifications that remain undetected at the level of global brain network emerge when the analysis focuses on restricted brain sub-networks. In line with previous findings, we found that AD impacted small world properties of the two brain sub-networks we identified, but closer investigation suggested that these modifications pointed in different directions: for one sub-network we found deterioration, for the other we found the possibility of improvement. Extending our analysis to other network parameters further revealed that complexity and diversity deteriorated in both networks. This new result enhances our understanding of the underlying pathology of AD in the human brain and suggests that in early AD, deterioration of brain networks may be accompanied by other types of changes.

## ACKNOWLEDGEMENT

We acknowledge the grants that contributed to the collection of OASIS data: P50 AG05681, P01 AG03991, R01 AG021910, P50 MH071616, U24 RR021382, R01 MH56584. Majnu John’s work was supported in part by grants from the National Institute of Mental Health for an Advanced Center for Intervention and Services Research (P30 MH090590) and a Center for Intervention Development and Applied Research (P50 MH080173). None of the authors has a financial conflict of interests.

---

**SUPPLEMENTARY FIGURE 1: Sparsity levels differ in AD patients and control groups for the same correlational thresholds.** Data of AD group are shown in red, data of control group are in blue. The two curves were obtained based on AD patient and control data, respectively, by computing sparsity (proportion of edges out of the total number of possible edges) of graphs constructed by using as correlation threshold all correlation values between 0 and 1 in steps of 0.001. One sparsity level corresponds to two different correlation coefficients for patient and control groups, and conversely, one correlation coefficient value corresponds to two different sparsity levels (dashed lines). Sparsity level of 5% corresponded to a correlational threshold of 0.5715 in patients and 0.5225 in controls; sparsity level of 34% corresponded to a correlational threshold of 0.3765 in patients and 0.3515 in controls.

